# Improving the Reliability and Quality of Nextflow Pipelines with nf-test

**DOI:** 10.1101/2024.05.25.595877

**Authors:** Lukas Forer, Sebastian Schönherr

**Affiliations:** Institute of Genetic Epidemiology, Medical University of Innsbruck, Innsbruck, Austria

**Keywords:** Nextflow, Pipeline Testing, Test Automation

## Abstract

The workflow management system Nextflow builds together with the nf-core community an essential ecosystem in Bioinformatics. However, ensuring the correctness and reliability of large and complex pipelines is challenging, since a unified and automated unit-style testing framework specific to Nextflow is still missing. To provide this crucial component to the community, we developed the testing framework nf-test. It introduces a modular approach that enables pipeline developers to test individual process blocks, workflow patterns and entire pipelines in insolation. nf-test is based on a similar syntax as Nextflow DSL 2 and provides unique features such as snapshot testing and smart testing to save resources by testing only changed modules. We show on different pipelines that these improvements minimize development time, reduce test execution time by up to 80% and enhance software quality by identifying bugs and issues early. Already adopted by dozens of pipelines, nf-test improves the robustness and reliability in pipeline development.

## INTRODUCTION

Due to the large amounts of biological data generated from various sources within genomics, proteomics, or metabolomics, the disciplines of Bioinformatics and Computational Biology turned into a big data science [1]. It includes both processing large datasets as well as applying complex workflows to analyze, filter and transform them to uncover complex biology relationships. Nextflow [2] has emerged as a powerful and flexible platform for building scalable and reproducible computational pipelines in the field of Bioinformatics. In conjunction with nf-core [3], a community-driven initiative dedicated to developing and maintaining best practices pipelines, a rich ecosystem has evolved [4]. However, as pipeline complexity grows, ensuring their correctness and reliability becomes a critical challenge, especially when incorporating new features without disrupting existing ones. While testing is a critical aspect of scientific software development [5], it remains underused in scientific software [6]. Thus, maintenance requires substantial time and effort to prove manually that the pipeline continues to produce scientifically valid results.

Automated testing is the process of evaluating and verifying that a software product does what it is supposed to do [7]. It is essential in scientific pipeline development to confirm the pipeline’s functionality and to ensure accurate data processing and analysis [8, 9]. It further involves defining test objectives, selecting test datasets, creating test cases, executing the pipeline with these test cases and verifying testing results [10]. Many Bioinformatics pipelines suffer from the so-called oracle problem, which means that they often handle large input and output data and implement complex algorithms without a clear gold standard [11]. This complexity makes the process of writing test cases complicated and time-intensive., A comprehensive and effective testing strategy for the pipeline’s functionality needs to cover different test levels and is divided into unit testing, integration testing and end-to-end testing. Unit testing plays a crucial role in software testing by verifying the correctness of individual units or components of code. Integration testing is important to ensure that different processes interacting within a larger workflow are working together as expected. Finally, end-to-end testing involves testing the entire pipeline from start to finish as a user would run it.

Despite efforts from existing solutions to automate end-to-end testing of Nextflow pipelines [12], a unified and robust unit-style testing framework specific to large and complex Nextflow pipelines is still lacking. This limits the efficient and automated validation of their functionality and makes it difficult for developers to guarantee the accuracy of their workflows, which could lead to errors and artifacts in data analysis and interpretation. This issue is even more relevant when considering the clinical utility of such pipelines. Moreover, long running execution times hinder developers from rerunning tests promptly, thereby limiting their productivity.

Here we present nf-test, a testing framework designed to address these challenges within the context of Nextflow pipelines. nf-test provides a domain specific language with similar syntax as Nextflow DSL 2 to describe the expected behavior and output data of a process or workflow. It introduces a modular approach that enables developers to isolate and validate individual process blocks, workflow patterns, and even entire pipelines. This modularity does not only simplify the debugging process but also encourages iterative development and code reuse. Moreover, we introduce snapshot testing and provide several optimization strategies to make testing data-intensive pipelines more efficient. This all helps pipeline developers to catch issues early in the development cycle and enables a robust and agile development process that results in more reliable pipelines. Serving as the new standard testing framework for nf-core [3], nf-test emerges as an essential tool for pipeline developers in the field of Bioinformatics. It is freely available and extensive documentation is provided on the website.

## RESULTS

### nf-test framework

nf-test is implemented as a command-line program, shares the same requirements as Nextflow and is compatible with Linux or macOS. We adapted well established testing concepts from software and web-development to Nextflow pipeline testing. The software offers a wide range of project-specific configuration options and can be easily installed on Continuous Integration (CI) platforms by using the provided installation script. Instructions, user guides and examples are available at https://www.nf-test.com.

### Unit testing in nf-test

Within the context of a Nextflow pipeline, unit testing involves testing a single process, workflow or function in isolation. We developed a Domain Specific Language (DSL) based on Groovy that offers methods and keywords to describe the expected behavior of any Nextflow unit. A project typically consists of different test suites, with one test suite per test subject (e.g., process, workflow, or pipeline). Each test suite contains one or more test cases to describe the expected behavior of the test subject. A test case is defined with the “test” keyword followed by two distinct blocks: (1) the “when” block, which set the input parameters of the test subject, and (2) the “then” block, which describes the expected output channels of the test subject when executed with the input parameters defined in the “when” block. Typically, the “then” block mainly contains assertions to check assumptions, such as the content of an output channel or files. Several functions are provided for writing assertions and simplifying Nextflow channel testing. Additionally, the ‘then’ block accepts any Groovy script and allows the import of third-party Java libraries.

This modularity also enables to write integration tests for a sub-workflow to ensure that the processes work together as expected. Thus, testing is consistently conducted using the same syntax and concepts throughout the entire Nextflow project. Together, these different levels of testing provide a comprehensive and effective test strategy of the pipeline’s functionality ensuring that all components work together as intended (see **Figure 1**).

**Figure 1:**
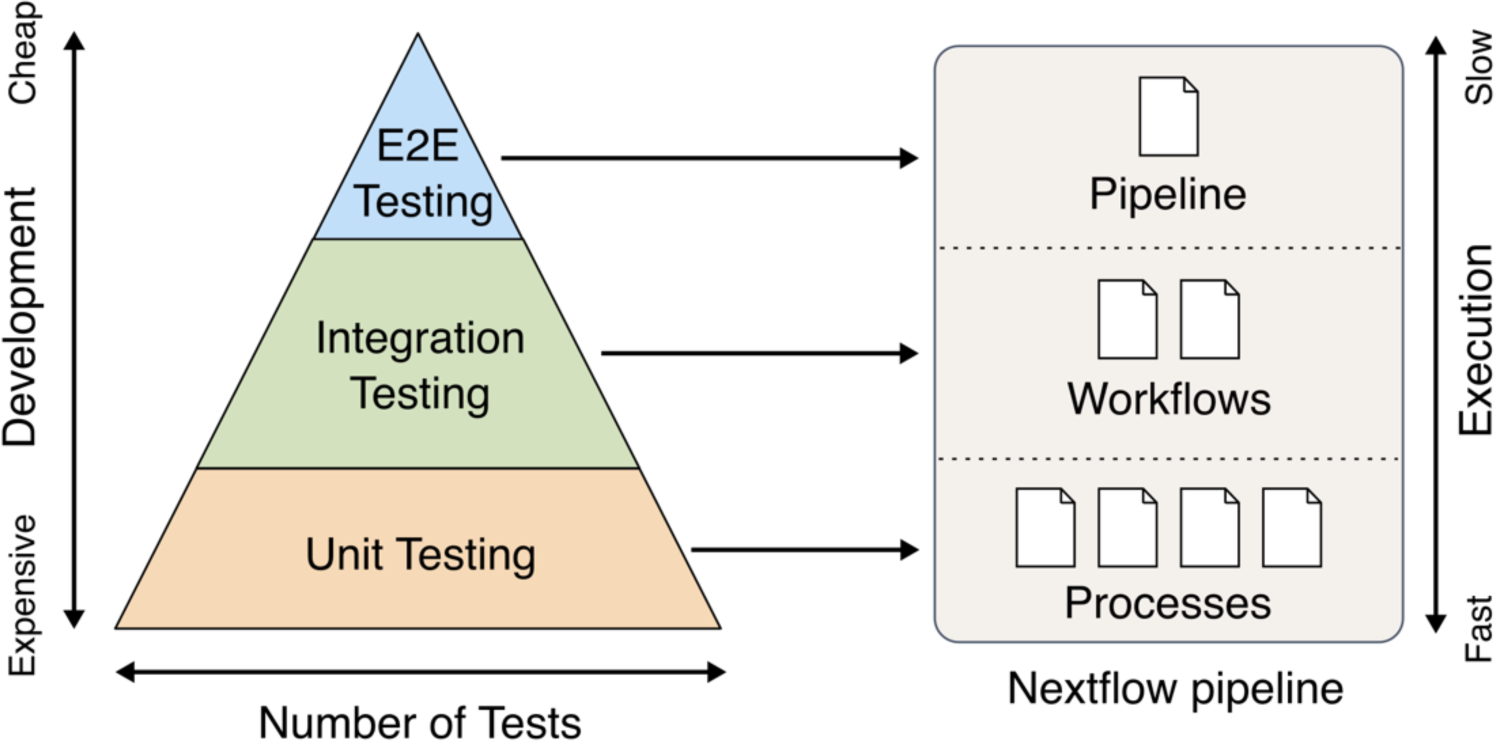
Overview of different test strategies. A compressive and efficient test strategy for a Nextflow project includes unit, integration and end-to-end testing.

### Testcase execution

One or more DSL files serve as an input for nf-test, which automatically executes the entire suite of tests. For each test, the runner automatically creates a Nextflow driver script that (a) initiates the Nextflow unit with parameters defined in the “when” block, (b) executes the unit and (c) serializes all output channels. Subsequently, it parses the content of the output channels and evaluates the assertions defined in the “then” block to verify if the output aligns with the expected behavior. As processes are executed in parallel, Nextflow channels emit output values in a random order. Deterministic assertions are made possible through sorting of the Nextflow channel tuples. The sorting is performed automatically by nf-test prior to the launch of the “then” closure. Finally, the test results are aggregated and reported in various formats (e.g., junit, xml, TAP (http://testanything.org/) or csv), allowing for processing by third-party reporting tools (see **Figure 2a**).

**Figure 2:**
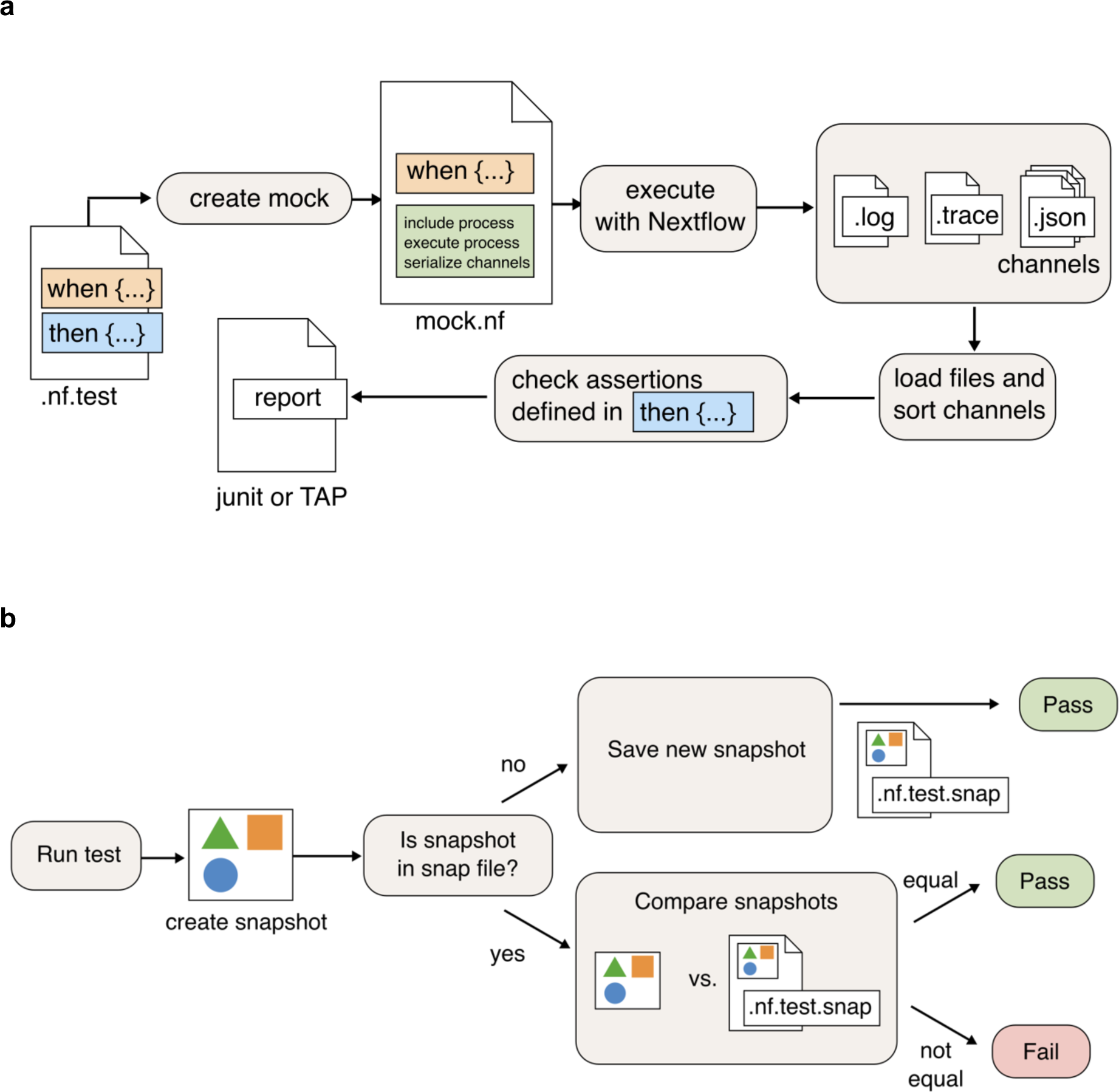
Architecture of the implemented test framework. **(a)** nf-test generates Nextflow scripts for tests: (i) initializes Nextflow unit with ‘when’ block parameters, (ii) executes the unit, serializes output channels, (iii) parses channel content, evaluates ‘then’ block assertions for output validation. Test results are aggregated and reported in multiple formats. **(b)** nf-test runs a test, creates and compares output objects against reference snapshot files stored with the tests. A test fails if the snapshots don’t match, indicating either unexpected changes or the need to update the reference snapshot to match new outputs.

### Smart testing and parallelization

nf-test provides a strategic approach called *smart testing* to optimize test execution time through minimizing the number of executed tests. We implemented a graph based approach inspired by firewall testing [13, 14] to select and prioritize tests based on their relevance to recent modifications. This enables nf-test to identify and execute tests for the impacted files and avoids redundantly retesting of unaffected components. As most Nextflow pipelines take advantage of a version control system, we have integrated Git (https://git-scm.com) support to automatically detect changes in the local working tree. Thus, it is possible to test changes between commits or branches, ensuring continuous validation of pipeline modifications on each commit or between releases. Parallelization is achieved through the implementation of *test list sharding* [15]. This technique divides the test suite into smaller subsets and distributes them across multiple parallel execution environments or cores. To optimize test distribution, we provide two distinct strategies: (a) simple chunking algorithm that splits the test list in chunks and (b) a round-robin approach for equitable allocation.

### Snapshot testing

We adapted the main idea of snapshot testing and applied it to the output of pipelines. Snapshot testing is a technique commonly utilized in web development to prevent unexpected changes in a user interface [16]. nf-test captures a snapshot of output channels or any other provided objects. These snapshots are then compared against reference snapshot files that are stored alongside the respective tests. If the two snapshots do not match, the test will fail (see **Figure 2b**). If the change is unexpected, the user can fix the bug detected by the test. Otherwise, the reference snapshot needs to be updated to the new output by rerun nf-test with a special option (‘--update-snaphot’). All snapshot files are created automatically in the background and are optimized for large files and complex objects.

### Extensions for Bioinformatics

nf-test provides a plugin system to reuse code snippets to save development time, to make test code cleaner and to enhance maintainability. The plugin system, based on Groovy, is well-documented to encourage users to create and share their own extensions. For example, in most Bioinformatics files formats file determinism is not always guaranteed, since timestamps or input filenames may prevent files to be byte-identically. In this cases md5 sums cannot be used and validating the dynamic output content can be time-intensive. We provide plugins for VCF, FASTA and CSV files to check the number of variants and samples as well as specific genotypes for a given sample.

### Resource saving through smart testing

First, we evaluated smart testing and its impact on execution time optimizations using the nf-core/fetchngs pipeline. This pipeline implemented 50 test cases for 17 components. The dependency graph illustrates the connections and dependencies between modules, workflows and test cases. Given that each component includes at least one test case, the pipeline achieves 100% coverage (see **Figure S1**). The total execution time for running all tests is 1,122 seconds. As hypothesized, we observed that pipeline end-to-end tests show the slowest performance (see **Table 1**). We simulated different modifications to assess the impact of changes, varying the number of modified files and different types of changes: (a) changes the logic of the module itself and (b) changes of the module interface (e.g. adding a new input channel). The results show that smart testing saves between 46% - 80% of the execution time by minimizing the number of executed testcases (see **Table 2**). We reviewed the test results with a full run of the entire test suite and confirmed that all changes are detected by our approach.

**Table 1:** Number of testcases in the nf-core/fetchngs pipeline 1.12.0. Each of the 17 components has at least one testcase. Total testcases 50. Total Execution time: 1,122 sec.

|  | Tests suites | Tests cases | Execution Time |  |
| --- | --- | --- | --- | --- |
|  |  |  | Mean (sec) | Total (sec) |
| Functions | 3 | 14 | 2.6 | 36.5 |
| Pipelines | 1 | 1 | 512.1 | 512.1 |
| Modules/Processes | 10 | 13 | 11.2 | 145.7 |
| Workflows | 15 | 22 | 19.4 | 427.8 |
| Total | 29 | 50 | - | 1,122.0 |

**Table 2:** Time and resource saving for different modifications of the nf-core/fetchngs. We simulated different typical modifications and measured the execution time using nf-tests optimization strategy. Time saving is calculated based on the execution time of a full run of 1,122 sec.

| Modification | Executed test cases | Execution Time |  |  |
| --- | --- | --- | --- | --- |
|  |  | Mean (sec) | Total (sec) | Saving |
| changed module sra_to_samplesheet | 10 | 22.3 | 222.5 | 80.2% |
| changed modules sra_to_samplesheet multiqc_mappings_config | 11 | 52.7 | 579.4 | 48.4% |
| update of a nf-core module: utils_nfcore_pipeline | 23 | 14.4 | 332.1 | 70.4% |
| changed main workflow main.nf | 1 | 596.3 | 596.3 | 46.8% |

Second, we analyzed the last 500 commits within the nf-core/modules project, spanning from October 26, 2023 to February 23, 2024. At the time of writing this paper, nf-core/modules contains 1,150 modules and 56 workflows as well as more than 800 testcases were implemented by the community. As expected, most of the commits and pull requests (PRs) affect only one module. In such cases, the time savings are significantly higher, since it only requires testing the relevant unit tests and potential integration tests of workflows. nf-test was able to accurately identify the specific test cases responsible for the committed changes and required integration tests (see **Figure 3a**). Around 30% of the commits needed the execution of more than 25 test cases, with most of these commits being refactoring or restructuring tasks (see **Figure 3b**). The total number of executed testcases could be decreased from 238,205 to 1,600. nf-test parsed and analyzed 1,560 unique files in under 1 second to construct the dependency graph.

**Figure 3:**
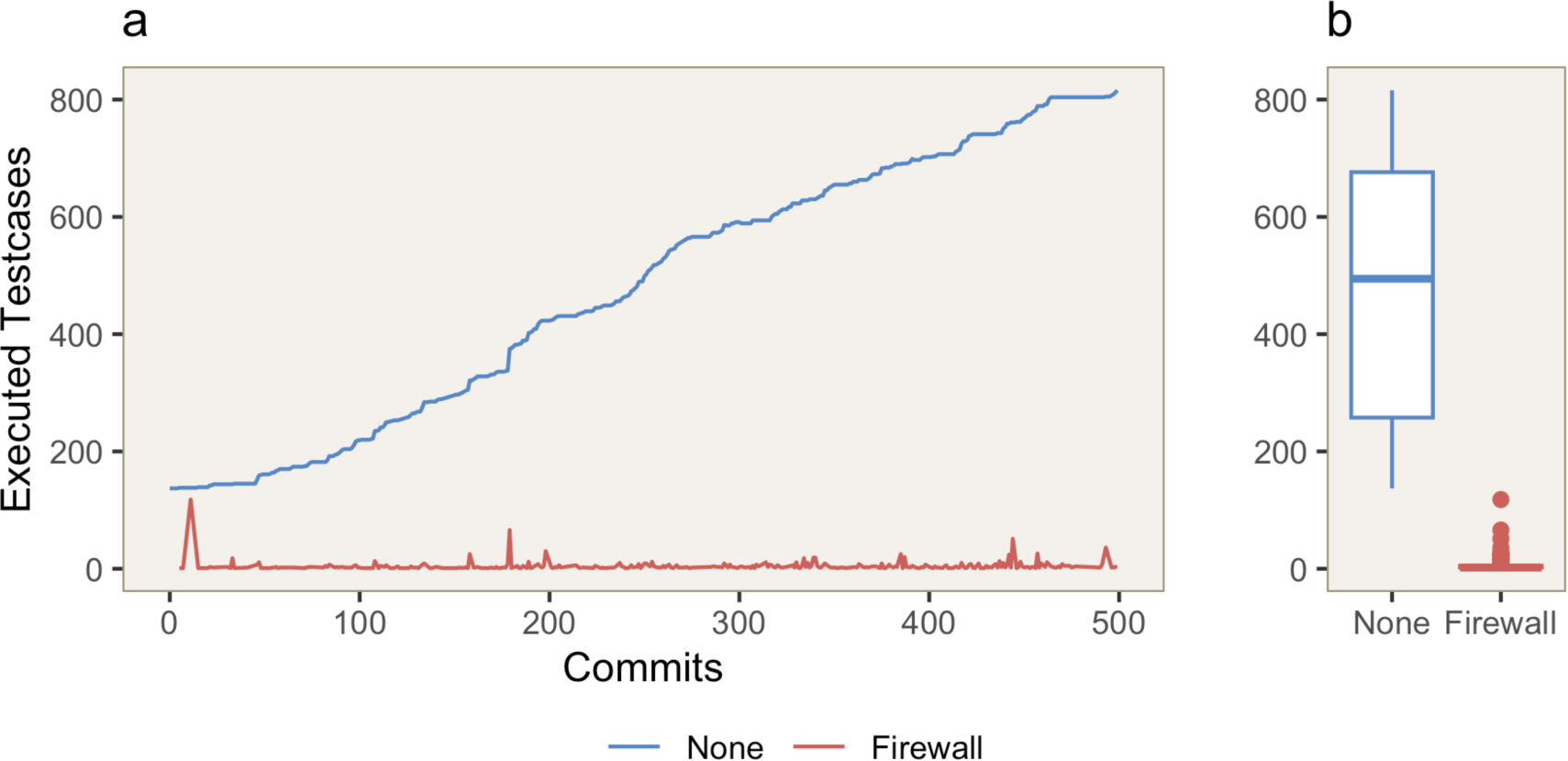
Last 500 commits of the nf-core/modules projects between 2023/10/26 and 2024/02/23. (A) The blue line represents the number of test cases that would be executed without any optimization strategy. The red line depicts the number of executed test cases using the implemented Firewall strategy. (B) Boxplot of the number of the executed testcases per commit.

### Execution time reduction through parallelization

We evaluated the efficiency and achieved speed up of parallelization on the example of nf-gwas [17]. It is a pipeline to run genome-wide association studies and consists of multiple long running end-to-end tests (see **Figure S2** for the dependency graph). Initially, executing the 49 test cases on a single machine required 1,718 seconds. The execution time could be reduced to 487 seconds by distributing the workload across five machines (Speed-Up: 3.5). The default strategy distributes tests by name, which may result in unbalanced execution times, especially when outliers are present (see **Table 3**). Using a round-robin approach further reduced the time to 333 seconds, resulting in a speed-up of 5.2.

**Table 3:** Time and speed-up for different sharding strategies of the nf-gwas pipeline.

| Strategy | Shards | Median (sec) | Total Time | Speed Up |
| --- | --- | --- | --- | --- |
| none | 5 | 368 +/-164 | 487 sec | 3.52 |
| Round-Robin | 5 | 314 +/- 10 | 333 sec | 5.15 |

### Code reduction through snapshot testing

We analyzed how snapshot-testing in nf-test could help to improve quality and maintainability by minimizing the effort of writing manual assertions. Writing manual assertions for each output item can be time-intensive and error-prone, especially in pipelines with extensive outputs. For example, the nf-gwas pipeline generates 5 output files for a single phenotype. Writing a simple regression test involves creating an assertion for each file to verify its existence and another to ensure its content matches expectations. This manual process implies generating an MD5 hash for each file and using it in the corresponding assert block, resulting in 15 lines of code for three phenotypes. Additionally, this testcase must be updated whenever the pipeline generates additional output files, requiring manual synchronization. Snapshot testing streamlines this process by replacing 15 lines of code with a single line. nf-test automatically creates the md5 sum for each of these files on the first run and provides commands to update to reference snapshot.

We adapted the nf-gwas pipeline and simulated various software updates for REGENIE: (a) changes in the output file format (renamed column “LOG10P” to “PVALUE”), (b) a bug in association detection (where 6 variants are no longer genome-wide significant), and (c) modifications to default parameters. Change (a) will break the pipeline and is easy to detect without testcases by running the pipeline with test data. However, changes (b) and (c) lead to incorrect output results without breaking the pipeline. Thus, these are the worst-case scenarios for a pipeline developer. The implemented test cases were able to detect all of these issues. For example, the unit test for REGENIE that checks if 116 variants are genome-wide significant fails, as there were only 110 variants found. Similarly, changes in default parameters (e.g., a more restricted MAF filter) lead to a different number of resulting variants and the test failed.

## DISCUSSION

Since its first release in October 2021, nf-test has been integrated in dozens of pipelines and was downloaded over 55,000 times. As the emerging standard testing framework in nf-core for both pipelines and provided modules, it shows a high level of acceptance, reflecting the community’s awareness of the importance of testing pipelines.

There were various efforts to test Nextflow pipelines in the past, including NFTest [12] and by the nf-core framework [3], which utilized pytest-workflow (https://pytest-workflow.readthedocs.io). However, these solutions are based on YAML files with predefined assertions, are limited to end-to-end testing, relied on external scripts to validate output files and did not support optimization strategies like parallelization and smart testing (see **Table 4**). nf-test addresses these gaps by offering a dedicated testing framework that simplifies the process of writing, executing, and analyzing tests for Nextflow pipelines. It offers a domain specific language (DSL) that follows a similar naming and philosophy as Nextflow DSL 2 and enables writing complex assertions that are often needed to validate the huge output of Bioinformatics analysis. As the DSL is based on Groovy, it can be extended by users themselves and they can rely on a rich ecosystem of Java/JVM libraries in the field of Bioinformatics (e.g., htsjdk and groovy-ngs). In addition, sharing domain specific assertions through plugins facilitate collaboration among users.

**Table 4:** Comparison of nf-test with similar approaches.

|  | **nf-test** | **NFTest** | **pytest-workflow** |
| --- | --- | --- | --- |
| **End to End Testing** | Yes | Yes | Yes |
| **Unit Testing** | Yes | Manual writing<br>Nextflow script | Manual writing<br>Nextflow script |
| **Snapshot testing** | Yes | Manual preparation of<br>expected output files | Manual preparation of<br>expected output files |
| **Optimized test**<br>**strategies** | Yes | No | No |
| **Dependency Analysis** | Yes | No | No |
| **Coverage reporting** | Yes | No | No |
| **Tags** | Yes | No | Yes |
| **Output Formats** | Junit, xml, TAP and csv | No | Junit, html |
| **Code generation** | config file and<br>for each unit | config file | Using nf-core tools |
| **Custom assertions** | Yes | Through external<br>third party scripts | No |
| **Bioinformatics**<br>**support** | Yes<br>(e.g. vcf, fasta, ...) | No | No |
| **Parallelization** | Sharding | No | No |
| **Modularity**<br>**and portability** | Each unit has its own test<br>files that can be<br>transferred | One test file for<br>whole project | Multiple files |
| **Writing Testcases** | Extendable DSL | YAML with<br>predefined structure | YAML with<br>predefined structure |
| **License** | MIT | GPL-2.0 | AGPL-3.0 |
| **Website** | https://www.nf-test.com | https://github.com/uclahs-cds/tool-NFTest | https://pytest-workflow.readthedocs.io |

We implemented a unit testing approach where all components of a Nextflow pipeline can be tested without the need for manually writing additional Nextflow workflows to execute a subprocess with test data. This modularity also facilitates writing integration tests for a sub-workflow to ensure that the processes work together as expected. All testing is conducted consistently across a project and possible introduced side effects are detected. This modularity does not only simplify the debugging process but also encourages iterative development and code reuse. Each module can be individually tested and seamlessly integrated into the final pipeline or workflow composition. Especially, when using large module libraries like the ones provided from nf-core/modules, integration tests are important to ensure that any third-party update doesn’t break its own pipeline.

In pipelines with extensive outputs, writing manual assertions for each output item can be time-intensive and error-prone. To mitigate this challenge, we introduced snapshot testing as a complementary approach. Instead of specifying individual assertions for each output, snapshots capture the state of the output channels or folders, including file names and hash values. These snapshots are automatically generated during the first run and nf-test is able to compare the current snapshot with the expected reference snapshot during every subsequent run. This makes regression testing more efficient and helps to catch regressions early in the development cycle. Moreover, it ensures that results remain reproducible and consistent across different runs of the same pipeline and after software updates. By evaluating the nf-gwas pipeline, we demonstrated how nf-test and snapshot-testing could enhance code quality and maintainability by minimizing the effort of writing manual assertions.

Regression testing involves retesting the pipeline after any code modification. Given that Bioinformatics pipelines process large input data and utilize complex algorithms, the execution of full regression tests could take hours. To save effort and time, nf-test only needs to retest those tests affected by the modification. We showed on the example of the nf-core/modules project, that the majority of changes and commits of such large projects affect only a specific set of modules and tests. Therefore, nf-test implements various approaches to detect minimal sets of tests that need to be rerun. This results in resource savings and faster development cycles (up to 80%).

When the number of tests in a pipeline is large and the execution time becomes long, test-list sharding can be used to split the tests across several machines. The experimental results indicated that this approach enables a significant performance gain of up to 80% in terms of reduction in execution time on fives machines. However, there is no guarantee of an optimal or fair split among resources since the splitting decision is not influenced by data from previous runs. The implemented round-robin strategy simply attempts to distribute the workload evenly. Nonetheless, the setup remains easy since no shared database, queuing system or orchestration instance is needed. Combined with integration of Git, nf-test enables the setup of Test-Driven Development (TDD) and Continuous Integration (CI) for Nextflow pipelines.

The implemented dependency analysis provides an overview of pipeline coverage and quantifies the testing effort. However, the current implementation has limitations, since it only reflects if at least one test case per unit exists, without indicating if all instructions or branches are covered by any test case. In future work, we aim to extend this approach and to include also metrics that reflect the complexity of a change set. Moreover, nf-test depends on local environments, which can make it less adaptable to certain infrastructures. For example, running test cases on different cloud providers is currently limited and will be addressed in future versions.

## METHODS

### Design and Implementation

nf-test is implemented in Java as a command-line program. We adapted well established testing concepts from software and web-development to Nextflow pipeline testing. nf-test is built upon a modular architecture and utilizes a plugin system, enabling effortless extension with new output formats, assertions, and optimization strategies. All source code is open source and freely available under the MIT-License.

### Smart Testing

#### Dependency Analysis

nf-test constructs a dependency graph that outlines the dependencies among all modules, workflows and test suites within a given project. In the graph, nodes represent individual modules, workflows and test suites, while edges represent the dependencies between them. The developed algorithm to create and discover these dependencies traverses through the entire project directory, identifies connections between different components (such as by parsing “include” statements), and maps them onto the graph structure. In **Figure 4**, an illustrative example showcases this concept. The nodes represent different processes (*M_1_*, *M_2_* and *M_3_*), workflows (*W_1_* and *W_2_*) as well as pipelines (*P_1_*) and are connected by edges denoting their dependencies (for example, *W_1_* depends on processes *M_1_* and *M_2_*). Test suites are connected to their testing subject (for instance, *T_M1_* is the test suite for *M_1_*). As a test suite could also be dependent on data or configuration files, it is possible to (a) define a list of files that always trigger a full retest (e.g. Dockerfile), and (b) define the assets of test cases which are automatically added to the dependency graph (e.g., input data). This directed graph offers insights into the underlying architecture of the pipeline, facilitating a comprehensive understanding of its dependencies and interactions.

**Figure 4:**
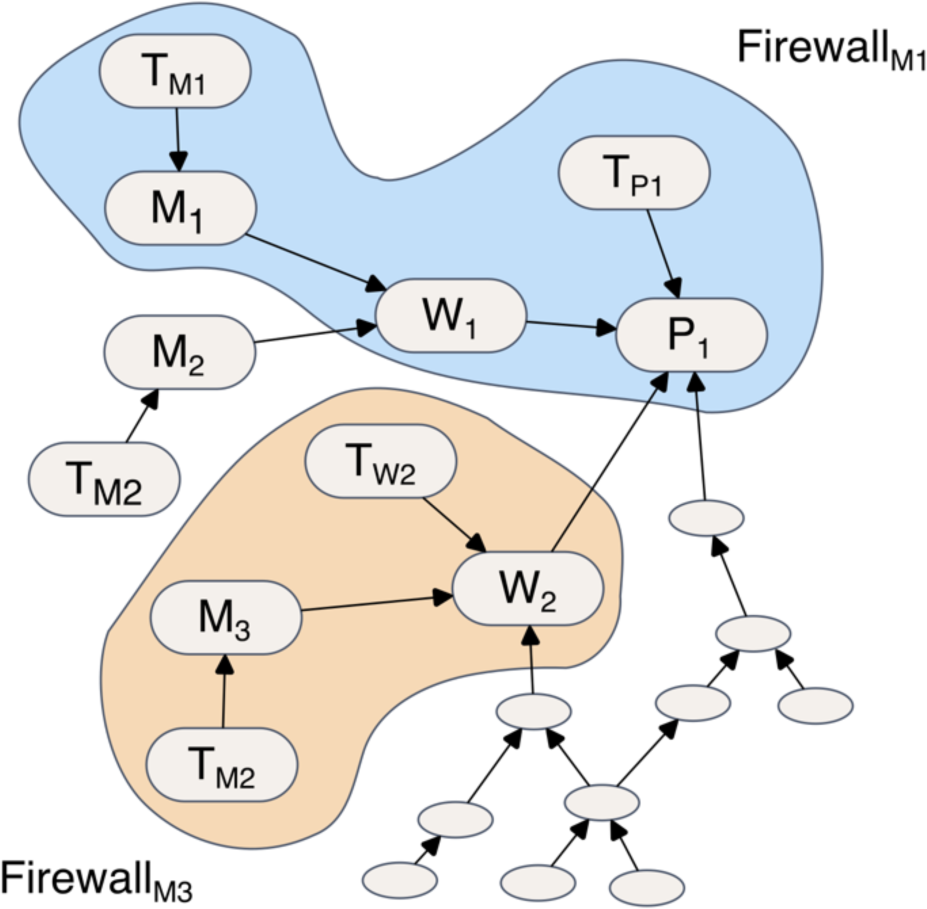
Example of a dependency graph and firewalls. The figure illustrates the dependencies of tests (*T_x_*), modules/processes (*M_x_*), workflows (*W_x_*), and pipelines (*P_x_*) within a pipeline. Changes to module *M_1_* will affect only test cases inside Firewall *M_1_*. Firewall *M_3_* is more compact since Workflow *W_2_* contains a test case ensuring the integrity of *M_3_*. This approach avoids running expensive end-to-end tests for *P_1_*.

#### Optimizing Test Execution

The dependency graph is used to implement a cost-effective testing strategy called smart testing. The main idea is to use the dependency information to identify a minimal set of tests required to detect potential regressions for changed processes or workflows. This improves efficiency and effectiveness in test execution since only specific test suites need to be retested. Inspired by a concept proposed by Leung and White [13, 14], we implemented a strategy based on *firewalls*. nf-test identifies all nodes affected by the modified module and constructs a firewall around the nodes, containing only tests that are affected by the change. Only these tests need to be retested. For instance, in **Figure 4**, *Firewall_M1_* contains all nodes that have been detected for retesting if module *M_1_* is changed. Tests associated with *M_1_* are first added to the firewall, followed by the inclusion of integration tests for *W_1_*, because it depends on *M_1_*. Since *W_1_* has no direct tests, we trace all nodes that depend on *W_1_*. In this case *P_1_*. For *P_1,_* a testcase exists, which is added to the firewall and can also be reused to indirectly test *W1*. On the other hand, if *M_3_* is modified, the resulting firewall is smaller, since *W_2_* has its own test case and there is no need to add *T_P1_* to the firewall. This strategy enables us to refine the firewall by potentially excluding expensive end-to-end tests while maintaining accuracy and reducing testing costs.

#### Test Coverage Calculation

Test coverage is calculated by analyzing the dependency graph and by determining which components are directly or indirectly covered by a test. The percentage of covered components relative to the total number of components provides a measure of the test coverage, indicating how much of the functionality is validated by the tests. Additionally, coverage can be calculated for a firewall to obtain a metric of how safe or well-tested a given change set is.

### Parallelization due to Test List Sharding

We implemented test list sharding to split the tests across several machines [15]. The discovered tests are sorted by type and filenames to ensure a deterministic order across different machines. To optimize test distribution across *n* machines, we provide two distinct strategies: (a) simple chunking algorithm that splits the test list in *n* chunks and (b) a round-robin approach for equitable allocation (*‘--shard-strategy round-robin’*). For instance, splitting a suite into three shards for distributed processing could be achieved by running one of the three commands on each machine: ‘*nf-test--shard 1/3*’, ‘*nf-test--shard 2/3*’, and ‘*nf-test -- shard 3/3*’.

### Snapshot Testing

Snapshot testing is a technique commonly utilized in web development [16] and has been adapted by us to apply it to Nextflow pipelines. The naming and parameters are highly inspired by jest (https://jest.org). nf-test captures a snapshot of output channels or any other provided objects and subsequently compares them to reference snapshot files stored alongside the tests. The snapshot file is a JSON file that contains, for each snapshot, a serialized version of its content. When a file is added to a snapshot, its MD5 hash sum is automatically included instead of the file content itself. Furthermore, it is possible to save the MD5 sum of the complete content of a snapshot, enabling the storage of a compressed version of complex and large content. As snapshot files are simple text files, they can be easily checked into version control systems and support human-readable differences. On each test run, nf-test compares the actual snapshot with the reference snapshot. If the two snapshots do not match, the test will fail. If the change is unexpected, the user can fix the bug detected by the test. Otherwise, the reference snapshot needs to be updated to the new output of a process, workflow, pipeline, or function. In this case, the user can run nf-test with the option *‘--update-snapshot*’.

Furthermore, associated snapshot files are automatically included in the dependency graph to prompt test modifications when a snapshot changes. We implemented a special continuous-integration (CI) mode that can be activated with the ‘--cì flag. When CI mode is activated, nf-test will not update the snapshot and the test will fail.

### Evaluation and Validation

We evaluated nf-test with three publicly available Nextflow pipelines to demonstrate the implications and benefits of testing pipelines with nf-tests in the field of Bioinformatics. Firstly, we assessed nf-core/fetchngs (Version 1.12.0, https://github.com/nf-core/fetchngs) by simulating four specific changes through manual file modifications and running nf-test with the *‘--related-tests*’ option. Secondly, we implemented 39 testcases and Continuous Integration (CI) using GitHub actions for the nf-gwas pipeline (Version 1.05, [17]). We tested it by employing test-list sharding across five machines with the option*‘--shard i/5’*, evaluating both default and round-robin strategies for test case distribution. Speed-up was defined as the ratio between execution times with and without sharding. Lastly, we evaluated nf-core/modules (Commit ca199cf, https://github.com/nf-core/modules) by running nf-test on the last 500 commits, utilizing the *‘--changed-since HEAD^*’ flag to capture changes between consecutive versions. All analyses were conducted using nf-test 0.9.0-rc2 and the results were visualized using R 4.3.3 and ggplot2 3.5.0.

## CODE AVAILABILITY

nf-test and the documentation are available at https://www.nf-test.com. The source code is available at https://github.com/askimed/nf-test and is released under the MIT license.

## ACKNOWLEDGMENTS

We would like to express our gratitude to the nf-core community for their support in this project, including their contributions of test cases and valuable suggestions. Special thanks go to Sateesh Peri, Maxime Garcia, Edmund Miller, Nicolas Vannieuwkerke, Harshil Patel, Adam Talbot and all GitHub contributors.

## CONFLICT OF INTEREST STATEMENT

None declared.

## Notes

### Competing Interest Statement

The authors have declared no competing interest.

