## Supplemental Material for "Improving the Reliability and Quality of Nextflow Pipelines with nf-test"

Institute of Genetic Epidemiology

Medical University of Innsbruck

Schöpfstrasse 41

6020 Innsbruck, Austria

### Figures

**Figure S1: Dependency graph of nf-core/fetchngs.** Green rectangles represent Nextflow files, blue rectangles indicate test cases for these files and white rounded rectangles denote snapshots.

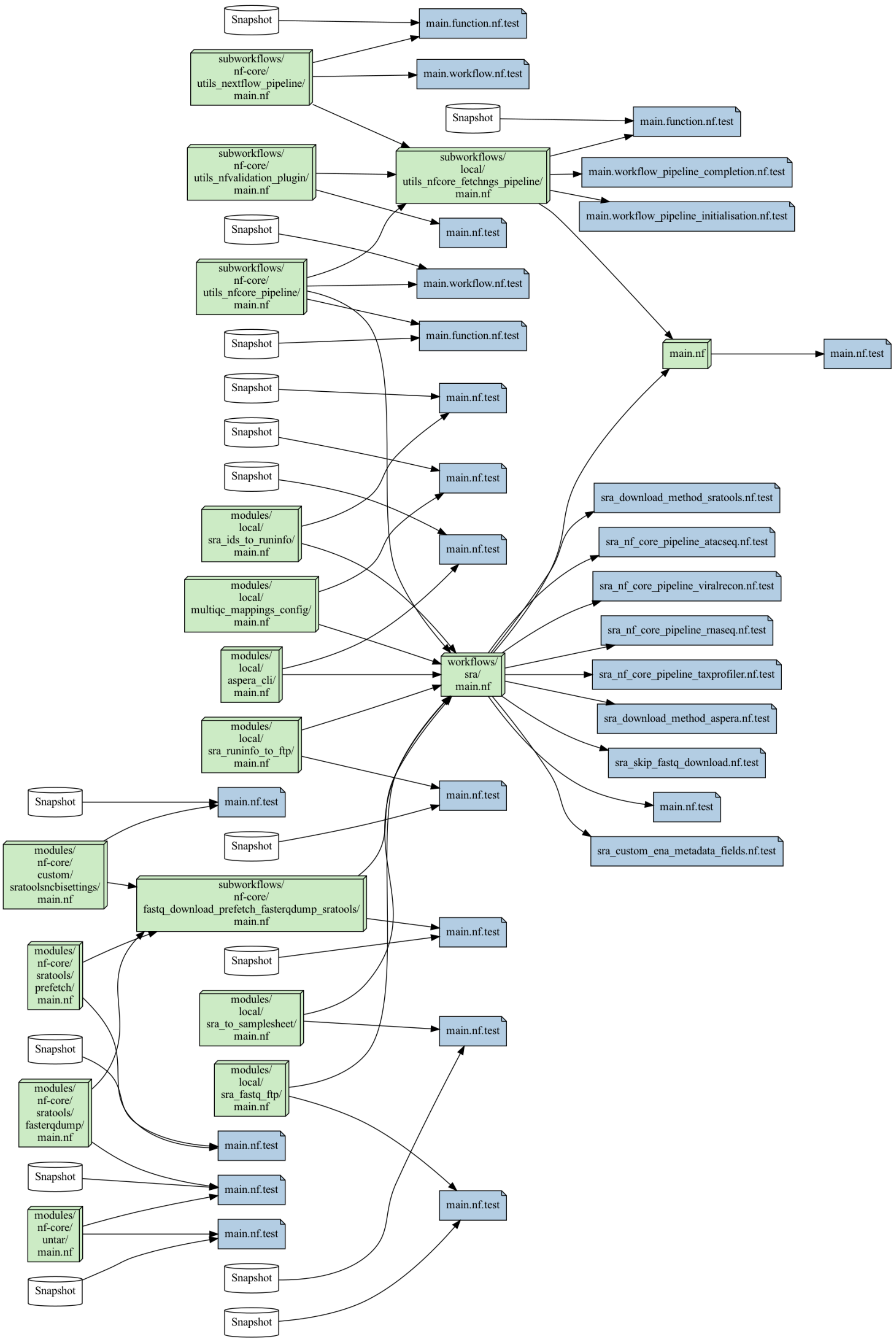

**Figure S2: Dependency graph of nf-gwas pipeline.** Green rectangles represent Nextflow files that have at least one test case, while red indicates those without a test case. Blue rectangles indicate test cases for these files and white rounded rectangles denote snapshots.

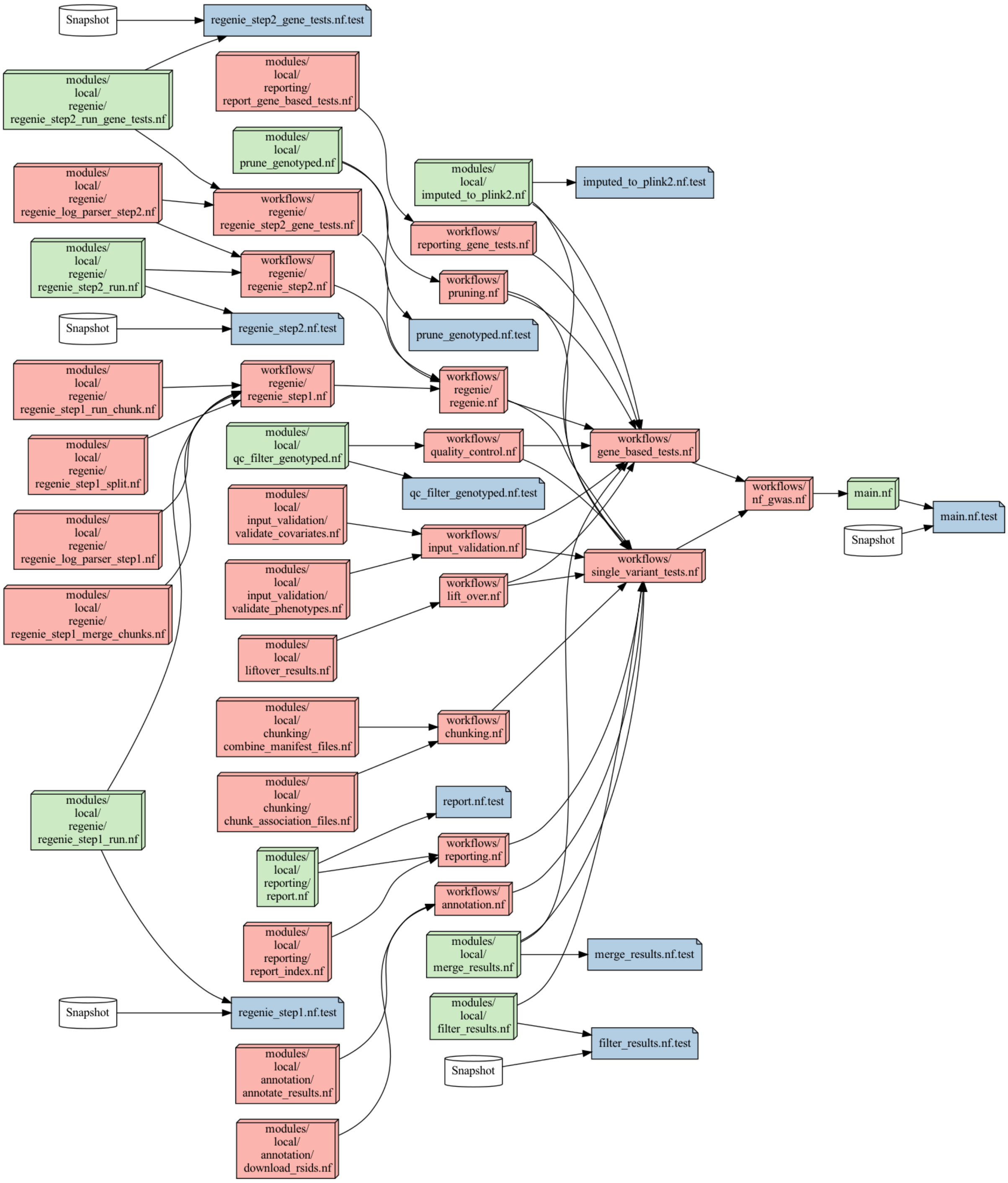
